# Are local dominance and inter-clade dynamics causally linked when one fossil clade displaces another?

**DOI:** 10.1101/2020.09.16.299750

**Authors:** Scott Lidgard, Emanuela Di Martino, Kamil Zágoršek, Lee Hsiang Liow

## Abstract

Disputing the supposition that ecological competition drives macroevolutionary patterns is now a familiar goal in many fossil biodiversity studies. But it is an elusive goal, hampered by patchy sampling, few assemblage-level comparative analyses, unverified ecological equivalence of clades and a dearth of appropriate statistical tools. We address these concerns with a fortified and vetted compilation of 40190 fossil species occurrences of cyclostome and cheilostome bryozoans, a canonical example of one taxonomically dominant clade being displaced by another. Dramatic increases in Cretaceous cheilostome genus diversification rates begin millions of years before cheilostomes overtake cyclostomes in local species proportions. Moreover, analyses of origination and extinction rates over 150 Myr suggest that inter-clade dynamics are causally linked to each other, but not to changing assemblage-level proportions.

**One Sentence Summary:** Global fossil diversification rates and local taxonomic dominance are not causally linked.

## Main Text

Time after time during life’s history, a major clade of organisms seems to be displaced by another with presumably similar ecological characteristics. This recurring pattern was once explained by facile assertions that competition gradually favored a better-adapted group over its rival (*1*). Now a new wave of studies is developing powerful process model frameworks aimed at narrowing the gaps between ecological interaction, changing taxonomic dominance, and deep time clade dynamics (*2–6*). In addition to more rigorous accounting for sampling and other biases, understanding how clade interaction and changing taxonomic dominance work depends on timing, rates and processes that encompass how organisms and their inclusive taxa interacted in the past (*6–10*).

The challenges to an integrated study of apparent clade displacement are formidable. In studies of fossil clade interactions, ecological equivalence is seldom well-established, and direct evidence of competition is rarely preserved (*2, 3*). Bryozoans are rare exceptions. Cyclostomes are closely allied to cheilostomes (*11*) and are comparable ecologically and physiologically, cooccurring on the same benthic substrates where access to space gives access to food (*11–14*). Their calcified colonies preserve competitive overgrowth interactions in fossil and modern assemblages (Fig. 1). Cheilostomes are routinely the overgrowth “victors” (*13*).

**Fig 1.**
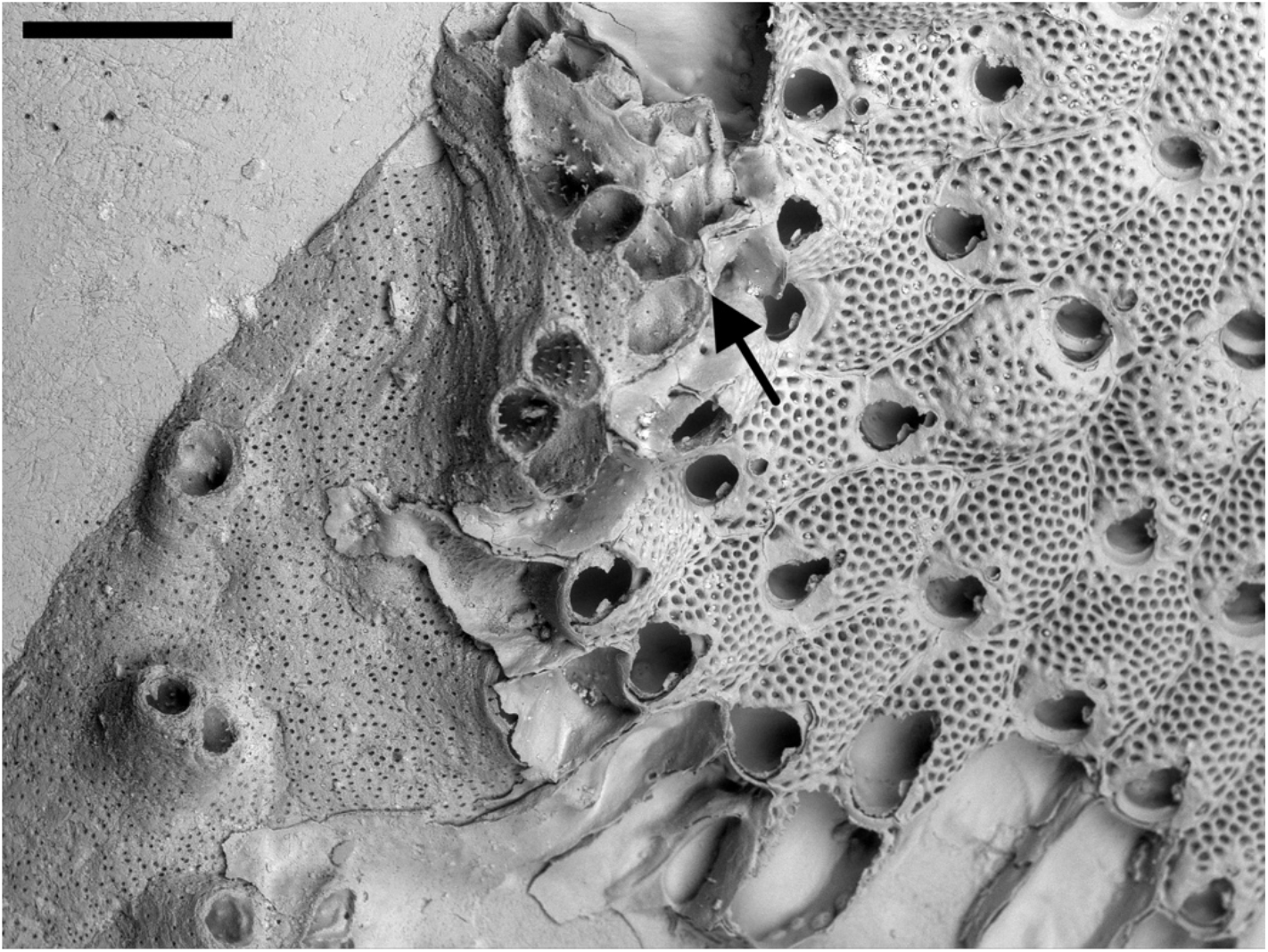
Competitive overgrowth. Encrusting cheilostome *Calyptotheca* at upper right overgrowing cyclostome *Annectocyma*, with partial “standoff” of raised colony margins (arrow). Recent, Cook Strait, New Zealand. Scale bar 500 μm.

Whether or not local ecological/species dominance is closely linked to diversification dynamics remains a largely unanswered question. Disappointingly few studies document changing species proportions of competing clades among individual assemblages (*12, 15, 16*). Clade displacement studies instead typically measure changes in global genus or family richness for each competing clade as proxies for changing patterns at lower scales (*7*). Insufficient synonymization may also bias diversity patterns (*17*). Appropriate statistical estimates of global diversity patterns are needed to account for sampling artefacts, requiring a sufficient set of occurrences through time, not least for grasping the uncertainty of diversification rates. The focus of clade displacement at a macroevolutionary scale is on changing rates of origination and extinction of taxa—are rates correlated between clades, or with changing local species proportions? Modeling these macroevolutionary processes provides a critical but seldom realized statistical basis for causal inferences (*2, 3, 6, 18*).

Here we present an integrated analysis designed to meet these challenges head-on for a canonical example of clade displacement (*1, 12, 19*). We compile data cataloguing cyclostome and cheilostome bryozoans over the past 150 Myr, identifying each genus/species, place of occurrence, and chronostratigraphic age (*20*). A greatly enlarged set of 40190 fossil species occurrences more than doubles the data in previous studies. We first analyze changing cyclostome-cheilostome species proportions within local fossil assemblages. Global genus richness patterns are then compared through 33 geologic stages, incorporating taxonomic revisions due to synonymizing and accounting for heterogenous sampling. We employ two disparate approaches (*2, 21*) to estimate the underlying genus origination and extinction rates while modeling sampling rates. Finally, we investigate potential correlative and causal connections of rates within and between clades and changing local proportions using linear stochastic differential equations (*20, 22*).

Species proportions in local assemblages are highly variable, but smooth splines fitted to the assemblage data (*20*) support an inference that the average within-assemblage proportion of cheilostome species started to exceed that of cyclostomes by ~75 Mya in the Campanian (Fig. 2A, species counts in fig. S1, sensitivity analysis in fig. S2). Estimates suggest a “crossover” to higher global genus richness of cheilostomes just after the Campanian (Fig. 2B). Genus estimates using the Jolly-Seber model (*2*) are robust to an alternative richness estimation (*21*) using PyRate (fig. S4). Raw non-synomymized data are comparable to synonymized data when range-through genus richness is visualized (fig. S3).

**Fig. 2.**
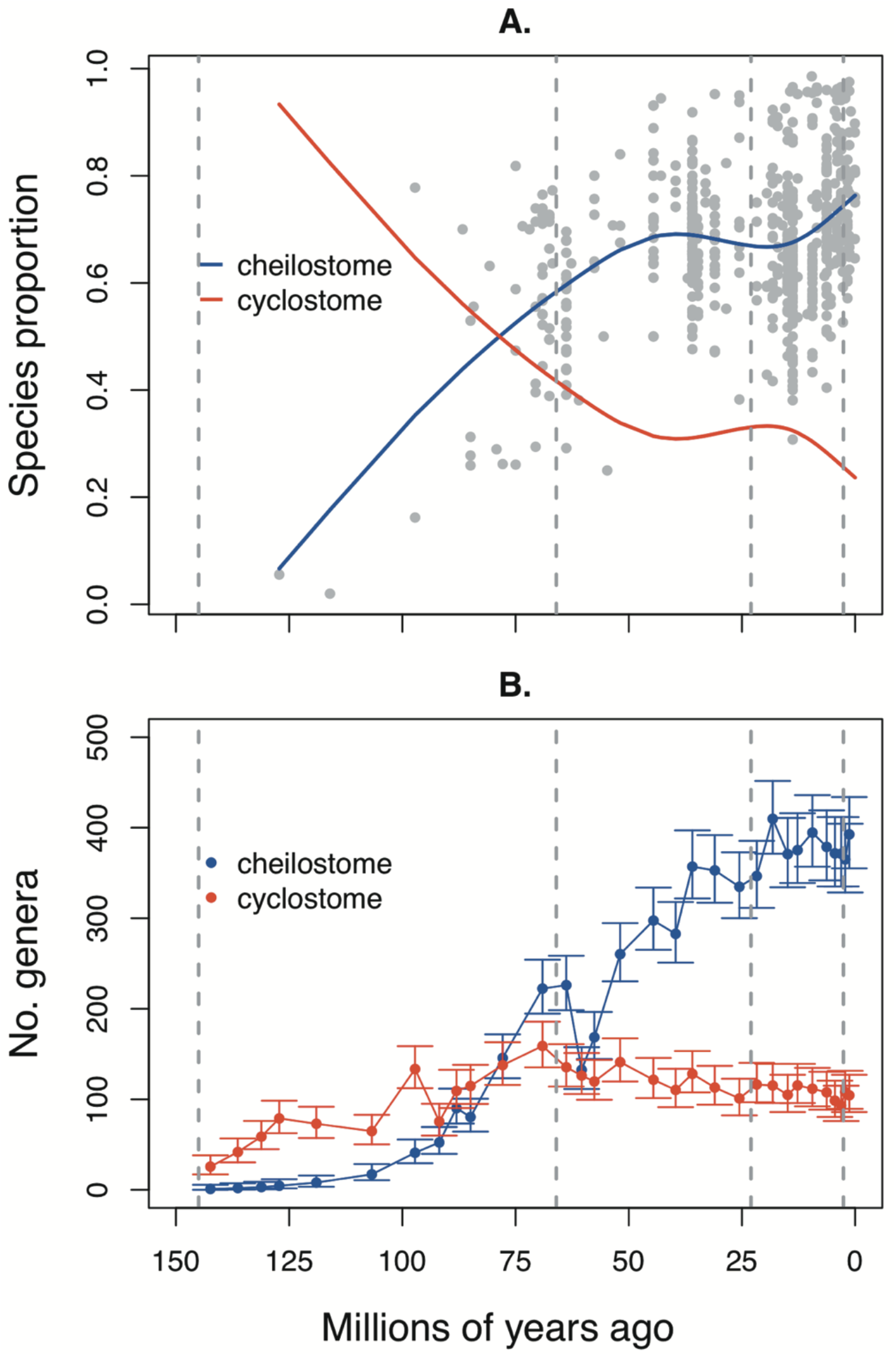
Changing taxonomic dominance at species and genus levels. A. Grey dots are within-assemblage proportions of cheilostome species. Only assemblages including both cyclostomes and cheilostomes and at least 15 species are shown (alternate species minimums in fig. S2). The blue line is a spline fitted to cheilostome proportions; the red line is the cyclostome complement. B. Genus richness estimated using a Jolly-Seber model and synonymized occurrences, with 95% confidence intervals (*20*). Vertical dashed lines from left to right delimit the Cretaceous, Paleogene, Neogene, and Pleistocene.

Despite fewer Mesozoic than Cenozoic data, analyses show clearly that cheilostome genus origination rates significantly exceeded cyclostome ones during the Cretaceous (Fig. 3A, S6A), and cheilostome extinction rates are mostly lower (Fig. 3B, S6B). During many Cretaceous and Cenozoic stages, higher cheilostome origination rates contrast sharply with cyclostomes, as do net differences between origination and extinction. These inferences are robust to alternative models (fig. S7-S10). Sampling rates are similar for cyclostomes and cheilostomes, even though estimated separately (Fig. 3C, S6C). Furthermore, Cretaceous cheilostome genus origination rates greatly surpassed cyclostome rates well before cheilostomes attained local species dominance (fig. S2, S6).

**Fig. 3.**
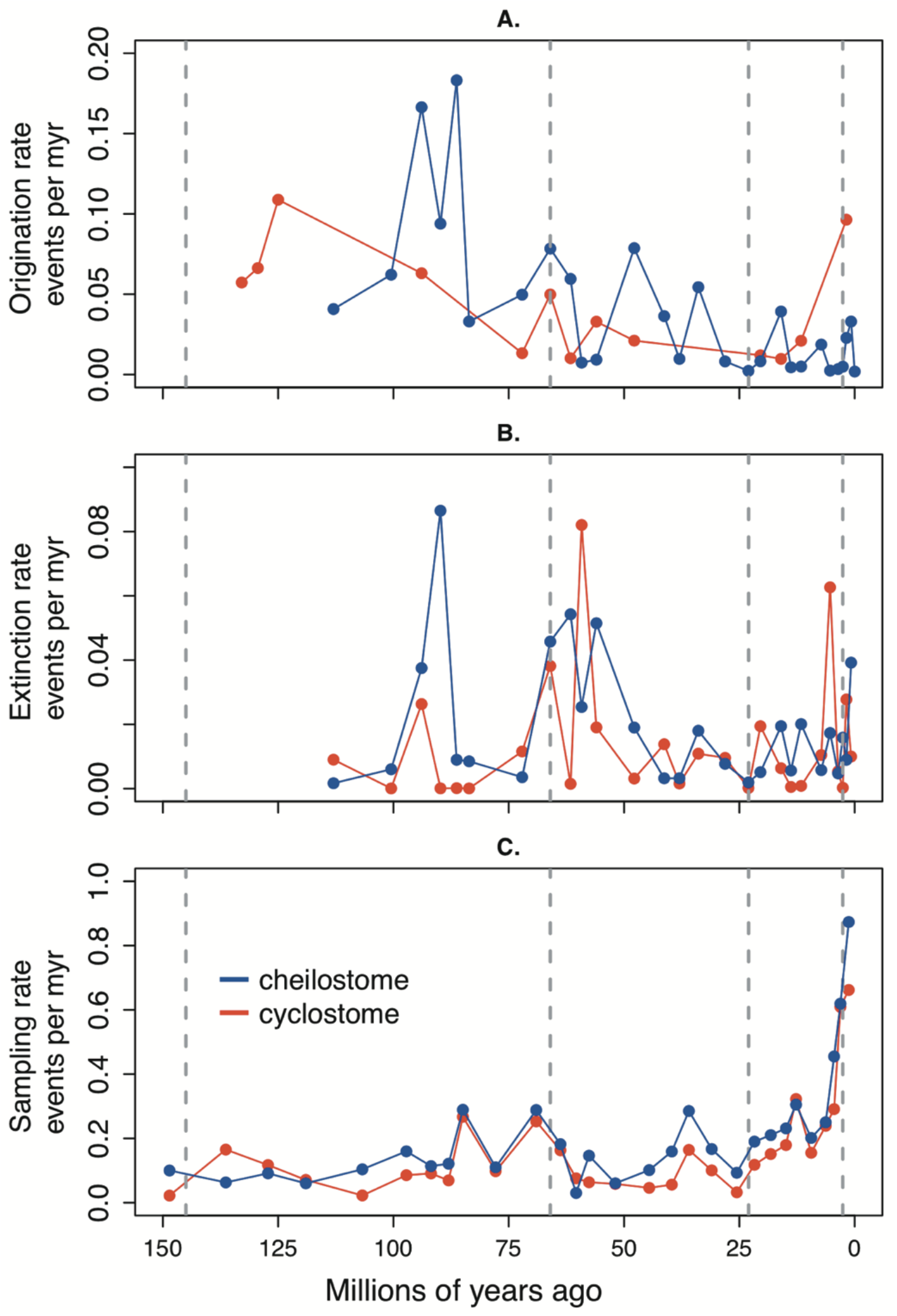
Genus origination, extinction and sampling rates. Each panel shows the estimates from fully time-varying Pradel seniority models run separately for cheilostome genera (blue) and cyclostome genera (red). Only relatively well-constrained estimates are plotted as dots (*20*, confidence intervals in fig. S5). Lines joining the estimates are for visual aid only. A. Instantaneous origination rates B. Instantaneous extinction rates. C. Sampling rates. Sampling rates are within-stage; origination and extinction rates are for stage boundaries.

We focus on statistically determined relationships among genus origination and extinction rates (*20*) in the upper portion of Table 1. From earlier prominent studies on clade displacement (*11–14, 19*), we hypothesized that cheilostome origination would dampen cyclostome origination (column 1, model B) and/or increase cyclostome extinction (column 3, model B), and that cheilostome extinction would facilitate cyclostome origination (column 4, model B). Surprisingly, cheilostome origination rates have no detectable relationship to cyclostome origination or extinction rates, given the weights of the null model (columns 1, 3, model A). These results suggest consideration of additional factors such as potential key innovations (*23, 24*) or environmental and ecological changes (*14, 25*), that could reveal relationships to pulses of cheilostome radiation. Between rates of cheilostome and cyclostome extinction, a feedback model has the highest model weight (column 2, model D). Intriguingly, we also detected feedback between cheilostome extinction and cyclostome origination (column 4, model D), rather than simple active displacement of cyclostomes by cheilostomes as indicated in past literature. The precise nature of feedback is uncertain, unsurprising as the time series themselves are uncertain and rather short. Considering the parameter estimates and Bayes factors between models (*20*), we can tentatively conclude that high cheilostome extinction rates could have led to a release for cyclostomes, facilitating a higher origination rate, although it is plausible that at the same time, high cyclostome origination rates could induce cheilostome extinction. In other words, one might speculate that rather than active displacement, one group’s partial disappearance could have resulted in another’s increased origination (*2*). Note that our models are process-based, analyzed for the entire time series rather than single events or short episodes, such as those spanning just two or three stages. While shorter-duration excursions in rates or proportions are “real,” they say little about statistical predictability in a Granger causal sense (*9, 20*).

**Table 1.** Causal links among diversification rates and within-assemblage proportions. Pairs of time series variables (columns) are evaluated using five different models of relationships (rows). Numbers in cells show the percent support for different models (*20*, parameter estimates in Table S1, S2). Grey cells indicate the best model. Abbreviations: cheilostome (Cheil) and cyclostome (Cycl) genus origination (orig) and extinction (ext) rates; proportions of cyclostome species to combined species in faunal assemblages (Proportion).

|  |  | Time Series |  |  |  |
| --- | --- | --- | --- | --- | --- |
|  |  | 1. Cheil orig<br>2. Cycl orig | 1. Cheil ext<br>2. Cycl ext | 1. Cheil orig<br>2. Cycl ext | 1. Cheil ext<br>2. Cycl orig |
| Models | A. No relation between time series | 41.8 | 11.4 | 49.2 | 24.1 |
|  | B. 1st time series drives 2nd | 10.5 | 22.9 | 11.4 | 9.8 |
|  | C. 2nd time series drives 1st | 20.4 | 13.4 | 14.5 | 24.4 |
|  | D. Temporal feedback between time series | 17.9 | 44 | 14.6 | 31.2 |
|  | E. Correlation between time series | 9.4 | 8.3 | 10.2 | 10.5 |
|  |  | 1. Cheil orig<br>2. Proportion | 1. Cheil ext<br>2. Proportion | 1. Cycl orig<br>2. Proportion | 1. Cycl ext<br>2. Proportion |
| Models | A. No relation between time series | 34.9 | 73 | 44.4 | 38.2 |
|  | B. 1st time series drives 2nd | 18.5 | 11.8 | 14.4 | 12.1 |
|  | C. 2nd time series drives 1st | 14.7 | 10.6 | 15 | 16.9 |
|  | D. Temporal feedback<br>between time series | 25.2 | 0 | 18.4 | 24.9 |
|  | E. Correlation between<br>time series | 6.6 | 4.6 | 7.8 | 7.9 |

Finally, in the lower part of Table 1, we examine relationships between genus origination and extinction rates and local species proportions. We hypothesized that global cheilostome and/or cyclostome origination and/or extinction rates could be driven by temporal changes in local faunal proportions. Instead, we find no relationship between any of the origination and extinction rates and species proportions represented in local assemblages (model A). This defies the notion that diversification rates are driven by local representation. Changing local species proportions and global genus rates do not even share the same dynamics, given the uniformly low weights for time series correlation (model E).

Looking ahead, greater resolution would be possible with more extensive sampling from the paleotropics and southern hemisphere (*26, 27*). A more efficient approach for all such fossil biodiversity studies could utilize Natural Language Processing methods to survey publications that are not yet incorporated into existing compilations (*28, 29*, fig. S5 and S7). Comparative modeling could also incorporate additional factors alongside competition (*30, 31*). Our analyses and new process model frameworks suggest a re-evaluation of the warrant for “bottom-up” competition in bryozoans and other well-known examples of fossil clade displacement.

## Acknowledgments

We thank D. Silvestro for advice on PyRate, T. Reitan for advice on causal analyses, and C. Schuette for help compiling and validating data.

## Author contributions

S.L. and L.H.L. designed the study, analyzed the data, and wrote the manuscript. S.L. collected and prepared the data, assisted by L.H.L., E.D.M., C.S., and K.Z. All authors contributed to editing.

## Data and materials availability

All data and code for analyses are available online at Dryad.

## Supplementary Materials

Materials and Methods

Figures S1-S10

Tables S1-S2

External Databases: see Dryad link

References (*1–46*)

