## Supplementary material for "Are local dominance and inter-clade dynamics causally linked when one fossil clade displaces another?": SI

**This file includes:**

**Data and materials availability:** All data and code for analyses are available online at Dryad.

### 1 Fossil bryozoan data sets

This study's primary data on fossil bryozoan genus and species richness includes three basic elements: identity of each genus/species, place of occurrence (location/region, see below), and chronostratigraphic age of occurrence. We begin with a bibliography of over 10,000 scientific references on bryozoans that had been compiled over many years and stored using the software *Zotero* (Roy Rosenzweig Center for History and New Media 2019). The bibliography includes references from previous analyses of cheilostome and cyclostome genus/species richness (Lidgard et al. 1993; McKinney et al. 1998, 2001). We filter the bibliography manually by tagging references whose titles (and some abstracts) suggest that their contents are likely to include lists of species from specific fossil localities or from circumscribed regions (e.g., Vienna Basin or Austria), or introductions of genus names. We prioritize such lists and genus introductions, rather than references detailing one or a few species, or mere mentions of species names in studies that do not focus on occurrences. The majority of relevant references are already stored as pdf files, and we retrieve most (but not all) others from online publication portals, or by scanning library text copies into pdf format and using optical character recognition software. We examine pdf files of tagged references, retaining those that include the three basic data elements in one of the two databases that are used for analyses (see below). Retained genus/species names are all identified as members of a calcified post-Paleozoic bryozoan clade, either Cyclostomata or Cheilostomata. In comparison to these other groups, ctenostomate and freshwater phylactolaemate bryozoans represent only a small component of modern bryozoan biodiversity. Because they are also uncalcified and their fossil occurrences are both rare and erratic, we exclude fossil ctenostome and phylactolaemate references and specific occurrences in genus/species lists. Fossil cheilostomes first appear in the latest Jurassic. We restrict the geologic time interval for reported fossil bryozoan occurrences under analysis to roughly 150 Mya through Holocene, thus excluding occurrences older than the Tithonian. We impose an arbitrary historical cutoff for references with publication dates older than 1920, yielding a set of literature spanning a century. For publications older than this date, especially from the mid- to late 19th century, determining locations of occurrence and their geographic extent, reliable geologic ages (particularly as disused regional stage names), and relative certainty of taxonomic determinations (and synonymizations) is exceedingly difficult. This cutoff may artificially deflate taxonomic richness for genus/species not reported for particular time intervals in subsequent works, but is intended as a compromise weighed against introduction of other sources of uncertainty, including correct recognition and taxonomic assignment (see below). While ours is the largest compilation of published Cretaceous to Recent bryozoan species occurrences to date, it is not all-inclusive. Indeed, all such fossil compilations are at best representative samples used to estimate true underlying patterns.

Primary data was extracted manually (but see fig. S5 and S7) from retained pdf files and stored in two databases (i.e., collections), *FosLocal* and *Age-Only*, facilitating analyses at local and global levels (Lidgard et al. 1993; McKinney et al. 1998; McKinney and Taylor 2001).

**FosLocal.** This database contains cheilostomes and cyclostomes reported from a single locality and stratigraphic level and identified to species level in the overwhelming majority of instances. If a publication reported species from more than one stratigraphic interval or horizon at a given locality, each one is retained as a separate entry. Thus, from a single reference, one entry may comprise just a single species, while another may comprise several or many tens of species. The geographic area of a locality varies considerably among references, but is intended to correspond roughly to a local fossil assemblage for purposes of within-assemblage species richness comparisons. A fossil assemblage is usually understood as "any group of fossils from a suitably restricted stratigraphic interval and geographic locality" (Fagerstrom 1964, 1198). Such assemblages can vary in the extent of pre-burial and post-burial transport, mixing and alteration of the original living community. Here we consider only members of the calcified bryozoan faunal community preserved in the fossil assemblage. Each locality is coded by the author's designated name or a similar abbreviation and originally assigned geologic stage (local or regional stage name, period or fraction thereof). We consult online chronostratigraphic literature and regional stratigraphic resources, both historical and modern, in an attempt to validate uncertain locality, formation, and age assignments (see also below).

The FosLocal database is used in our analyses of within-assemblage fossil species proportions of cyclostomes and cheilostomes. In many genus/species lists, authors include taxa that they were unable to determine precisely to genus or species rank. Indeed, species counts from older literature are likely to underestimate actual richness. Advances in scanning electron microscopy, morphometrics, and molecular genetic methods of species discrimination over the past four decades have shown that morphologically cryptic species do occur among traditionally recognized bryozoan species. Where publications list some species as questionable determinations, we have retained these taxa, treating them according to their relevance for particular analyses. For instance, species identified as similar to a known name, such as *Callopora* aff. *dumerilii*, were retained as the known name, *Callopora dumerilii.* Species designated as multiples, such as *Callopora* spp., were counted only once, as *Callopora* sp., since we do not know how many different morphospecies were actually recognized. Species only designated as belonging to a higher taxonomic rank or identifiable group, such as Calloporidae sp. or Cribrimorph sp., were assigned to either cyclostomes or cheilostomes for within-assemblage counts (e.g., Cheilostome sp.). This database contains 1695 locality entries and 36155 genus/species occurrences.

The FosLocal database is combined with the Age-Only database for our analyses of fossil genus richness of cyclostomes, cheilostomes, and all calcified bryozoans combined. Species designated simply as belonging to a higher rank or identifiable group, such as Calloporidae sp. or Cyclostomata sp., are excluded from global genus richness analyses (see sections 2.3 to 2.7).

**Age-Only.** This database includes genus/species names that do not necessarily come from a single locality, but are constrained temporally to a single geologic stage (or, in some cases, consecutive stages) and constrained geographically to a localized region (i.e., state, country, or depositional basin). For instance, Labracherie (1971) summarizes lists of bryozoans known from successive Eocene epochs in the North Aquitaine sedimentary basin. Also included are references that introduce genus names or describe genera and species of an entire family or higher taxonomic rank together with occurrences. Both of these types of references may include multiple localities, but do not correspond to full lists of species found at a single locality. Examples include the introduction of *Quadriscutella* gen. nov. from the Tertiary and Recent of Victoria and South Australia by Bock and Cook (1993), and the classic study of Cretaceous Pelmatoporine cribrimorph cheilostomes by Larwood (1962). This database contains 308 locality or regional entries and 4418 genus/species occurrences.

The Age-Only database is combined with the FosLocal database for our analyses of fossil genus richness of cyclostomes, cheilostomes, and all calcified bryozoans combined.

1.1 <u>Geologic age standardization</u>

We assign updated estimates of geologic age to each entry in our databases. For most entries we rely on the stratigraphic placement given by the author, but necessarily revise those that refer only to regional geologic stage names, obsolete or generally disused stage names or lithostratigraphic units (including some 'formations'). This process involved consulting online and published literature sources for lithostratigraphic and chronostratigraphic verification. Representative sources include the GeoWhen Database (Rohde 2005), Geoscience Australia Stratigraphic Units Database (Geoscience Australia 2020), GEOLEX (U.S. Geological Survey 2020), publications revising or redefining regional stratigraphic units (Laga and Louye 2006, King 2016). For consistency, all data entries are then mapped onto corresponding stages given in version 2018/08 of the International Chronostratigraphic Chart of the International Commission on Stratigraphy (ICS, 2018). All analyses employ chronological ages from this version of the Chronostratigraphic Chart. Where original stage designations span more than one international stage, the combined successive stages are used to bracket an absolute age estimate. Where original stage designations referred to part of a stage (i.e., lower, upper), we adopt a convention of dividing the relevant international stage into equal halves to provide a chronological age bracket analogous to these divisions. In the FosLocal database, 459 of 1236 localities (37%) spanned more than one ICS stage while in the Age-Only database, 77 of 231 localities (33%) spanned more than one ICS stage.

1.2 <u>Taxonomic standardization</u>

Changes in taxonomic nomenclature of genus/species names are part of an ongoing process, one clearly reflected in a bibliography of references spanning a century. We use three authoritative sources of accepted names and previously applied synonymizations and invalidations (i.e., *nomen nudum*, *nomen oblitum*) to standardize genus names. These sources, the Working List of Genera and Subgenera for the Treatise on Invertebrate Paleontology (pers. comm. Dennis P. Gordon, 2019), World Register of Marine Species (WoRMS Editorial Board, 2020), www.bryozoa.net (Bock, 2020) are the most up-to date and thoroughly investigated ones available. Gordon's Working List contains accepted names of living and fossil cheilostome bryozoans. Bock's bryozoa.net treats all bryozoans and includes accepted names, known names whose status remains uncertain, synonyms, and invalidated names. These two databases and the WoRMS database are largely in agreement for accepted cheilostome names. The bryozoa.net database is larger than the others; it includes a number of previously published cheilostome and cyclostome genus and species names that have yet to be fully investigated. Both Gordon and Bock are editors for the Bryozoa section of the WoRMS database, which is relatively comprehensive for living taxa and also includes many fossil taxa and some synonyms. We incorporate minor updates to the list of valid names in the WoRMS database to include the most recent genus introductions that have not yet been uploaded to WoRMS. This validated list of living bryozoan genera is used to complement our fossil data for genus range-through analyses (see 2.3-2.7, below).

We compile an extensive set of 323 (cheilostome) + 125 (cyclostome) = 448 previously applied genus synonyms and invalidations from these sources and from the historical bryozoan literature. We filter all of the genus names in our two databases in order to compare the effect of genus-level nomenclatural updates on genus richness trends with and without synonymization into currently accepted taxonomic standards.

In all but one instance, we do not attempt to standardize species names that had resulted in synonymization of an author's original species name. In particular, synonymization to both a new genus and a new species epithet remains especially problematic. Complete research—likely including direct SEM examination of specimens—that would accomplish the latter kind of revision for all species would be an enormous long-term project, one simply beyond the scope of our study. One exceptional case is the disused genus "*Lepralia" sensu lato*, defined by Brown (1952) as being "used in a wide sense for encrusting Ascophora the generic affinities of which are not clearly understood." For species occurrences of *Lepralia*, we update genus and species designations to their modern equivalents, as most species historically assigned to this genus have been synonymized previously.

### 2 Detailed methods

#### 2.1 Within-assemblage species richness and proportions

For all localities in the FosLocal database, counts of cheilostome, cyclostome, and all calcified bryozoans combined are plotted in figures at the midpoints of their age ranges, using the International Chronostratigraphic Chart of the International Commission on Stratigraphy (fig. S1, N= 1689). These within-assemblage counts may include data from references that report only cheilostomes, only cyclostomes, or species of both groups. Visualization reveals overall trends in within-assemblage species richness over time, varying density of sampling among intervals, and several intervals in which some assemblages record exceptionally high species richness.

Proportions of cheilostome versus cyclostome species within assemblages are calculated using subsets of the localities in FosLocal database in which the author reported the presence of both cheilostomes and cyclostomes (N = 975 of the original 1689). In Fig. 2A, we estimate the timing of "cross-over" in higher species proportions from cyclostomes to cheilostomes by fitting separate cubic splines through the reported proportions of cheilostome and cyclostome species within assemblages, using the smooth.spline function in R (R Core Team 2019). These proportions, and hence the cubic splines, can be affected by the minimum number of species reported per assemblage. We hence analyze sensitivity of within-assemblage percentage trends to minimum species numbers per assemblage at three levels (fig. S2), retaining localities with at least 10 species (765 localities), 15 species (630 localities), and 20 species (531 localities). We also varied the degrees of freedom of the cubic spline from 3 to 13 to visualize the sensitivity of the "cross-over" point (fig. S2).

2.2 Genus range-through richness

Genus occurrences include all genus-level names recorded from both the FosLocal and Age-Only databases. Species originally identified only as members of a family or higher taxonomic level (i.e., Calloporidae sp.) are excluded. This filtered, combined FosLocal and Age-Only database is henceforth termed the combined database. Our approach differs from previous family or genus fossil bryozoan biodiversity compilations based only on reported first and last stratigraphic occurrences of taxa, and assumed continuity between those occurrences (e.g., Lidgard et al. 1993; Sepkoski et al. 2000; McKinney and Taylor 2001; Taylor and Waeschenbach 2015). We retained 694 synonymized cheilostome genera and 261 synonymized cyclostome genera in the interval between 150 Myr and the Holocene.

We tabulate range-through genus richness within each 2018/08 International Commission on Stratigraphy stage/age starting from the Tithonian (152.1 to 145.0 Mya), but merging the last three very short stage/ages into a single Holocene interval (i.e. Greenlandian, Northgrippian and Meghalayan are collapsed into one), such that there are a total of 33 unequal duration time intervals for our analyses. We first assign each observation to one of our time intervals using the mid-point of their age range. Additionally, we check if the genera represented in our combined fossil database are known to be extant, based on the World Register of Marine Species (WoRMS Editorial Board, 2020). If they are, we indicate that they are extant in our last time interval (Holocene), regardless of whether there is an observation of that genus in our combined database.

A genus is hence assumed to range from the time interval where it was first observed to the one where it was last observed (including the WoRMS "boosted" Holocene time interval). This implies that the genus cannot be extant before its first observation or last observation (see sections 2.4 onwards for how we relax this assumption). Range-through genus richness within each of the time intervals is simply tabulated as the number of genera directly observed, plus those inferred to have ranged through the time intervals where they are not observed but that are flanked by observations. We compare our range through genus richness with equivalent numbers tabulated in McKinney and Taylor (2001). As far as we are aware, it is the most recent publication in which stage by stage genus totals are presented (rather than plots alone). Even though McKinney and Taylor (2001) is an older compilation, subsequent reviews are based on it (e.g., Taylor and Waeschenbach 2015). We extracted data from Taylor and Waeschenbach 2015 using WebPlotDigitizer [https://automeris.io/WebPlotDigitizer/] and found that it almost exactly matches data in McKinney and Taylor 2001 (data not shown).

2.3 Genus range-through richness (synonym-corrected)

The tabulation of range-through richness here are performed in the same manner as described in 2.2, except that any original taxon names that are known to be synonyms (as described in Section 1.2; files cheilostome_SYNONYMS_MASTER_01jul20.xlsx and cyclostome_SYNONYMS_MASTER_01jul20.xlsx are available in the Dryad link) are substituted with the accepted names where available. 2887 lines of data representing 153 genera are substituted.

2.4 Genus richness estimation

While range-through genus tabulations can account for extant genera in some time intervals in which they are not sampled (i.e., inferred to be extant in certain time intervals because they are found both earlier and later time intervals), this basic form of tabulation does not utilize the information from the non-observation of genera. Additionally, confidence intervals for genus richness cannot be generated naturally in range-through tabulations. As noted above, the range-through approach assumes that a genus cannot have existed before its first observation or after its last observation, and that the only genera that can be unobserved are ones that have observations in two or more time intervals in the dataset. These are assumptions that will be relaxed here. Non-observations imply either true absence (i.e., a genus has not originated or has gone extinct) or the lack of sampling (i.e., a genus is extant but not sampled). To utilize such information in genus richness estimation, we use a Jolly-Seber model (Jolly 1965, Seber 1965), reviewed in Pollock et al. (1990). The Jolly-Seber model is an open population model in the capture-recapture literature (reviewed in King 2014) that estimates "population size" (i.e., genus richness in our data), "survival rates" and "birth numbers" (i.e. number of originations in our data). One of the most important limiting assumptions of this model for our application is that all genera have the same probability of being sampled. For instance, we expect very lightly calcified genera to be rarely preserved and very rarely sampled; hence our conclusions are likely reflecting preservation of moderately- to well-calcified genera. Other assumptions, including short sampling intervals relative to the time over survival is estimated, and independence of genera in the dataset have been explored and discussed extensively but are not thought to bias estimates (reviewed in Pollock et al. 1990, but see also Williams *et al.* 2002, Liow and Nichols 2010 and references therein). We implement the Jolly-Seber model using the JS.direct function in the openCR (Efford 2019) R package. To estimate 95% confidence intervals for the number of genera estimated, we assume a Poisson model of sampling.

For a comparison to another approach that explicitly models sampling probabilities but in a wholly different manner, we also ran PyRate analyses. Here, the preservation probability of each genus is assumed to change through its lifetime but estimation is conditioned on at least one observation of a genus (Silvestro et al. 2014a). We use the same genus occurrences (described at the beginning of section 2.2) and applied PyRate software (Silvestro et al. 2014b). A non-homogeneous Poisson process of preservation is assumed for our genera (the NHPP option), where we also specify a time-variable Poisson process for our time intervals (TPP option, Silvestro et al. 2019) and a gamma model of rate heterogeneity across genera (mG option). We increase the default number of iterations to 50000000, sampling every 5000th iteration. Convergence is checked in Tracer (Rambaut et al. 2018). We analyze cyclostome and cheilostome data separately for consistency with capture-recapture analyses. Note that estimates from PyRate are much "smoother" because the algorithm samples a point estimate from the temporal range of each genus occurrence as input into Bayesian analyses. To allow estimates of genus richness to be comparable with those from the Jolly-Seber model, we plot means from counts of genera in each international stage and 95% credibility intervals from 100000 runs. Posterior samples from PyRate are supplied in csv files in Dryad.

2.5. Genus origination and extinction estimation

For genus origination and extinction rates, we use the capture-recapture model called the Pradel seniority model (Pradel 1996) which combines forward and reverse-time modeling to examine both "survival" and "seniority," whose complements translate to extinction and origination for genus level data from the fossil record. This model has been used and described in e.g., Connolly and Miller (2001), Liow et al. (2008), Martins et al. (2018), and Sibert et al. (2018). The Pradel seniority framework is flexible in the sense that extinction, origination and sampling probabilities can be time-varying or may include covariates (e.g., cheilostomes and cyclostomes could be constrained to have different estimates in the same model). We first explore ten different models by using all the data in the combined dataset simultaneously (see R Markdown file CMR Markdown 01.07.2020. Rmd in Dryad) . However, we found that the best model by AIC criteria has time-varying sampling probabilities for cheilostomes and cyclostomes combined, and separate but constant extinction and origination probabilities for cheilostomes and cyclostomes separately. As we are interested in time-varying rates, we chose to estimate full models separately for cheilostomes and cyclostomes while accepting that estimates in some time intervals may not be well-constrained. We run such models for our combined dataset as well as a text-mined dataset (Kopperud et al. 2019), using the openCR.fit function while specifying "Pradelg" (Efford 2019).

The origination and extinction estimates from the Pradel model are transition probabilities across temporal boundaries. Sampling probabilities, however, are associated with the time interval in question. All estimated probabilities are transformed into instantaneous rates, as our 33 time intervals are unequal in duration, by assuming a Poisson model (see Liow et al. 2015). The probability of no event (i.e. origination, extinction or sampling) within a time interval is $P(no\ event)\ =\ e^{\lambda T}$, where $\lambda$ is the event rate and T is the time interval length. Probabilities that are 0 are not amendable to this log-transformation and are always associated with very large confidence intervals; hence, they are removed from further analyses.

As mentioned above, we use the posterior MCMC samples from PyRate analyses (described in the previous section) to present estimated origination and extinction rates for a comparison. We note there that we favor capture-recapture estimates for our analyses for several reasons. First, in the capture-recapture estimates, the uncertainty in origination and extinction rates stem directly from the distribution of genera known to be extant but otherwise unsampled. In PyRate, however, the uncertainty stems from the estimation of start and end points of taxon lifespans instead. The differences in uncertainties are in part reflected in their size differences, where Pyrate uncertainties are smaller but in our view less realistic (compare capture-recapture uncertainties with PyRate credibility intervals in fig. S4, S9, S10). PyRate also estimates origination and extinction rates given that taxa are sampled at least once, but capture recapture models relax this assumption, hence also contributing to larger (but more realistic) uncertainties. Last, PyRate models are designed to detect rate shifts, which is not our primary goal here.

2.6. Causal analyses using linear SDEs

We are interested in whether the taxon origination and extinction time series may be associated with or perhaps even casually linked to one another or to proportional changes in local assemblages. The statistical concept of Granger causality (Granger 1969) is employed across many disciplines as a probabilistic method for investigating *predictions* between time series (Wiederman and von Eye 2016). While it is common to test for correlations between time points in search for associations directly among time series data (e.g., papers using correlations or window shifted analyses in ACF) or to use first differenced data, we have shown that these approaches are prone to false inferences (see Liow et al. 2015 and review in Hannisdal and Liow 2018). Hence, we use a time series approach based on the work of Granger (1969) and Schweder (1970) to tackle causality. More specifically, we use linear stochastic differential equations (SDEs, see Øksendal 2010 for an overview and Reitan et al. 2012 for an application) to model Granger causality.

In brief, we use a system of linear SDEs to explicitly model temporal correlations and Granger causality among time series while relaxing the requirement of sampling at equidistant points, and embracing uncertainties associated with the time series estimates. In addition to comparing null, correlative and causal models, we also provide parameter estimates from each model, including estimates of the strengths of correlations and Granger causal relationships among time series. We implement this using the recently developed R package layzeranalyzer (Reitan and Liow 2019). We log transform the four extinction and origination rate time series to conform to the normality requirement in our linear SDE tool kit. We apply pairwise comparisons among the five time series rather than a multi-time series comparison, as our data series are too short for a full-scale analyses (compare Liow et al. 2015 and Reitan and Liow 2017 for consequence of a less acceptable pairwise analysis) and show the results of model selection in main text Table 1. We present parameter estimates (Tables S1 and S2) for the time-series pairs (Table 1) in which the null hypothesis is clearly rejected.

2.7 A comparison with text-mined data

The data we compile here are highly labor-intensive and require specialist knowledge in several aspects. We were curious as to whether an equivalent dataset mined using Natural Language Processing (NLP) tools would yield salient features of cheilostome and cyclostome diversification that were inferred using our combined database. The pipeline for mining such data was developed in an earlier publication, where range-through cheilostome genus richness was presented (Kopperud et al. 2019). We use the pdfs our bibliography (described in section 1) as a source of information and mine taxon-age candidate pairs (see Kopperud et al. 2019 for details). The mined data we present here are not synonym-corrected, but filtered for those that have more than 0.5 probability of being correct. This text-mined occurrence dataset (TMO henceforth) is at best only about one-third of the data volume compared with our combined dataset but it is not possible to directly compare the correspondence of data rows. We present genus range-through and Jolly-Seber estimates using TMO data as a comparison to estimates from the combined dataset.

### 3 Supplementary results and discussion

#### 3.1 Assemblage results

Species counts in assemblages do not exceed 50 species for either cheilostomes or cyclostomes at the start of our time series (>100 Million years ago, fig. S1). However, while cyclostome species numbers in assemblages remain relatively stable and low throughout, cheilostomes begin to contribute species richness to assemblages from about 100 million years ago (fig. S1). By plotting average proportions of cheilostomes and cyclostomes in each time interval and fitting splines to these averages, we see that the cheilostome and cyclostome splines "cross" during the latter part of the Campanian (~ 75 MA), no matter which data subset or however many degrees of freedom are used in spline-fitting (fig. S2). Our analyses reveal considerable differences in local species proportions relative to previous compilations (e.g., Lidgard et al. 1993). The "crossover" from higher cyclostome to cheilostome within-assemblage species proportions is estimated a few million years earlier, but average late Eocene cheilostome species richness is far greater, Oligocene declines more severe (with lowered cheilostome proportions), and more pronounced fluctuations in average richness occur between Neogene stages (Fig. 2A, S1, S2).

#### 3.2 Range-through tabulations

Simple range-through genus richness estimates put the total number of genera at the last time interval (Holocene) at about 470, with small fluctuations since the start of the Neogene, of which about 20% are cyclostomes and 80% are cheilostomes (fig. S3A). A comparison with the most comprehensive published compilation (McKinney and Taylor 2001) shows that our new compilation has contributed a substantial amount of new data, and changes due to synonymization of taxa in published literature. Considering only the past 100 Myr, we report 693 synonymized cheilostome genera and 238 synonymized cyclostome genera, versus 545 and 263, respectively, in McKinney and Taylor (2001). The end-Paleocene through Eocene cheilostome radiation proves to be massive, finally ~35% more in estimated genus richness than recognized previously. Synonymized cheilostome genera are >100 more numerous by end-Eocene. For cyclostomes, changes appear more modest. Note that for data around the KPg boundary and over the Cretaceous, our new compilation shows fewer range-through genera than McKinney and Taylor (2001). This is due in part to the synonymizations that have been introduced, but likely also due to our treatment of age data (fig. S3B).

3.3 Jolly-Seber and PyRate genus estimates

Genus richness estimates from PyRate and the Jolly-Seber (JS) model are similar in terms of both general temporal trends and mean estimates (fig. S4, using the same synonymized combined dataset). 95% confidence intervals from the JS model almost always overlap the means of the estimates from PyRate, but not the other way around. This is because the 95% credibility bands are in general very narrow (see section 2.4). The discrepancies between the cyclostome genus estimates are larger than those for cheilostomes: PyRate estimates are in general lower (closer to the lower bound, i.e., the range-through tabulations, see fig. S3). Genus estimates using a Jolly-Seber approach on the text-mined data for cheilostomes closely follow those using synonymized, manually collected data (fig. S5), whereas there are greater discrepancies for cyclostome genera that could not be reconciled even with a Jolly-Seber approach that takes into consideration incomplete sampling (fig. S5).

3.4 Pradel and PyRate origination and extinction estimates

In our main text, we showed plots comparing cheilostome and cyclostome origination and extinction rate estimates (Fig. 3A and B). For completeness, we also show the confidence intervals estimated in the Pradel model in fig. S6. There is substantial uncertainty reflecting the nature of the data. What is clear, as stated in the main text, is that cheilostome origination rates are in general higher than those of cyclostomes while extinction rates for the two clades are similar.

We also explored the use of text-mined data as a means of increasing the efficiency of gathering data for these types of analyses. Pradel model estimated cyclostome origination and extinction rates using text-mined data are higher than those for our manually collected data (fig. S7), even though the estimated genus richness (fig. S5) is lower. We could not estimate extinction and origination rate confidence intervals for many time intervals in which there were few observations of cyclostome genera in the text-mined data. Note, however, that the confidence intervals estimated from our manually collected cyclostome data overlap these point estimates without estimable confidence intervals. This overlap indicates that estimates from the sparse text-mined data still fell within the range of more confident estimates (fig. S7). Conversely, cheilostome origination and extinction estimates using text-mined data are lower than those for our manually collected data while confidence intervals largely overlap (fig. S8). In future, text-mined data for other groups (especially with more data) can be a beneficial compromise between manual effort and reliability of the diversification/richness estimates, if sampling is included as a parameter in the modelling.

Extinction and origination rates from PyRate analyses are not directly comparable with those from the Pradel model. The latter are transition rates from one time interval to the next, while those from PyRate plotted in figs. S9 and S10 are averages within time interval tabulated from MCMC (Monte Carlo Markov Chain) runs. The two classes of models also incorporate different assumptions. Despite these differences, many features of diversification processes are consistent between the two model estimates, even though the credibility intervals from PyRate are much smaller than the confidences intervals from the Pradel model. For instance, there is a substantial increase in origination rates for cheilostomes around 100 to 75 million years ago, giving us confidence that this predates the increase in assemblage representation of cheilostomes around 75 million years ago. Because we feel that PyRate gives credibility bands that may be too small, we chose to use estimates from the Pradel model with its larger confidence intervals as input for our causal analyses, which are designed to take account of patchy data.

We do not compare preservation (sampling) rates for the two models as they are very different in both assumptions and structure. More specifically, in PyRate, preservation rate is estimated as the expected number of fossil occurrences per sampled lineage per time while in the Pradel model, sampling rate is the probability that a lineage is preserved then sampled given that it is extant.

3.5 Causal analyses

Drawing upon previously published bryozoan studies, we hypothesized that cheilostome origination will dampen cyclostome origination or increase cyclostome extinction. Neither of these hypotheses are supported (main text upper Table 1, columns 1 and 3 ). Rather, cheilostome origination has no detectable relationship to cyclostome origination, or cyclostome extinction for that matter. This implies that the origination of cheilostomes has other underlying causes that are not related to assemblage level dynamics (main text lower Table 1, columns 1 and 3) or global diversification rates.

We also hypothesized that cheilostome extinction should be correlated with cyclostome extinction (main text upper Table 1 column 2, where model E should have been favored), as the two ecologically equivalent groups might be impacted by similar external drivers (which we did not model). Also, cheilostome extinction might "promote" cyclostome origination (main text upper Table 1 column 4, where model B should have been favored and where the parameter estimate for $\beta$ should have been positive). Again, neither of these hypotheses were supported by our data. Cheilostome extinction rates seem to have experienced feedback with cyclostome extinction rates (Table 1, column 2, model D. is favored). On examining the parameter estimates, we see that the nature of this mutual feedback is uncertain (Table S1). The point estimates of the $\beta$ 's in Model D (the feedback model) are positive, but the credibility bands span 0 and are quite "even" on both sides. On simplifying the model, to look at one causal arrow at a time (models B, and C. in Table S1), the estimates are more certainly positive. The Bayes factors between model D. and model A. and Model E. and A. (Table 1) are 3.85 and 5.3 respectively, indicating that there is substantial evidence that both Models A. and E. should be rejected. The uncertainty of the result is not surprising as the time series is highly uncertain and rather short (a longer or more densely parsed time series might improve resolution among models and their relative likelihoods). In other words, we can tentatively conclude that there is feedback between extinction rates and that greater extinction rates for cheilostomes lead to greater extinction rates for cyclostomes and also vice versa.

The uncertainty of the situation is similar for the relationship between cheilostome extinction and cyclostome origination rates, although clear rejecting our hypothesis. The two best models are the feedback model (Model D.) and the model in which cyclostome origination rates positively drive cheilostome extinction rates (Model C.). These two models (models C and D in main text Table 1 and Table S2) are not very different, but we can exclude the correlation model (using the rule of thumb of Bayes factor >2) and also the null model, if we are willing to sum over all three causal models to compare with the null model. However, just as in the previous case, the parameter estimates are very uncertain (Table S2), even though the point estimates are positive. This suggests but does not confirm that high cheilostome extinction rates could have led to a release for cyclostomes, facilitating a higher origination rate; extrapolating still further, when cyclostomes originated at a higher rate, cheilostomes were negatively impacted.

### 5 Supplementary figures (S1–S10)

**fig. S1. Raw data for faunal assemblages through time.** Each dot shows the actual species counts observed for the individual assemblages for A. cheilostomes (blue) and B. cyclostomes (red) separately. In A. light blue circles show assemblages where cyclostomes were not recorded while in B. pink ones show where cheilostomes were not recorded. Both of these latter sets were excluded from our analyses of within-assemblage proportions in the main text. Both groups experience a sharp decline in within-assemblage richness for the Paleocene stages. Cheilostomes increase spectacularly through Lutetian to Priabonian (~47.8-33.9 Mya), greatly outpacing cyclostomes. Both groups again decline through Rupelian and Chattian (~33.9-23.0 Mya), then increase with fluctuations through mid-Miocene (~14 Mya), after which cheilostomes show a much greater increase. Two outliers beyond the upper boundary of these plots are not shown: "Rugen, Oberes Mukronaten-Senon" (Voigt 1930; Lower Maastrichtian, 217 cheilostomes) and "Wilmington, N.C." (Canu and Basseler 1920; Priabonian, 158 cheilostomes and 56 cyclostomes). Noise is added to the ages of the assemblages to help visibility.

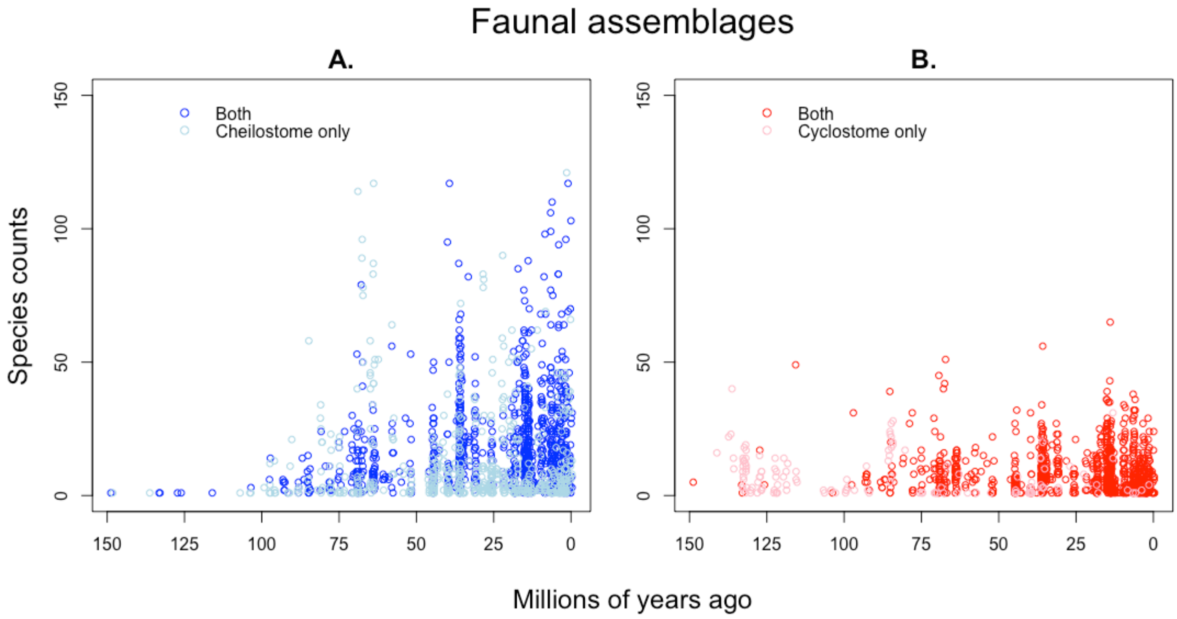

**fig. S2. Proportions of cheilostome and cyclostome species in assemblages through time.** Points are the means of proportions of cheilostome species (blue) and cyclostome species (red) in each international stage. Panels show variation depending on which assemblages are retained (e.g. N=>10 retains only assemblages with at least 10 species reported). Dashed lines show cubic splines with degrees of freedom increasing from 3 to 13 (dashes with more inflections have higher degrees of freedom), 2 steps at a time, for cheilostome (blue) or cyclostome proportions (red). Grey vertical dashes delimit the Cretaceous, Paleogene, Neogene, and Pleistocene.

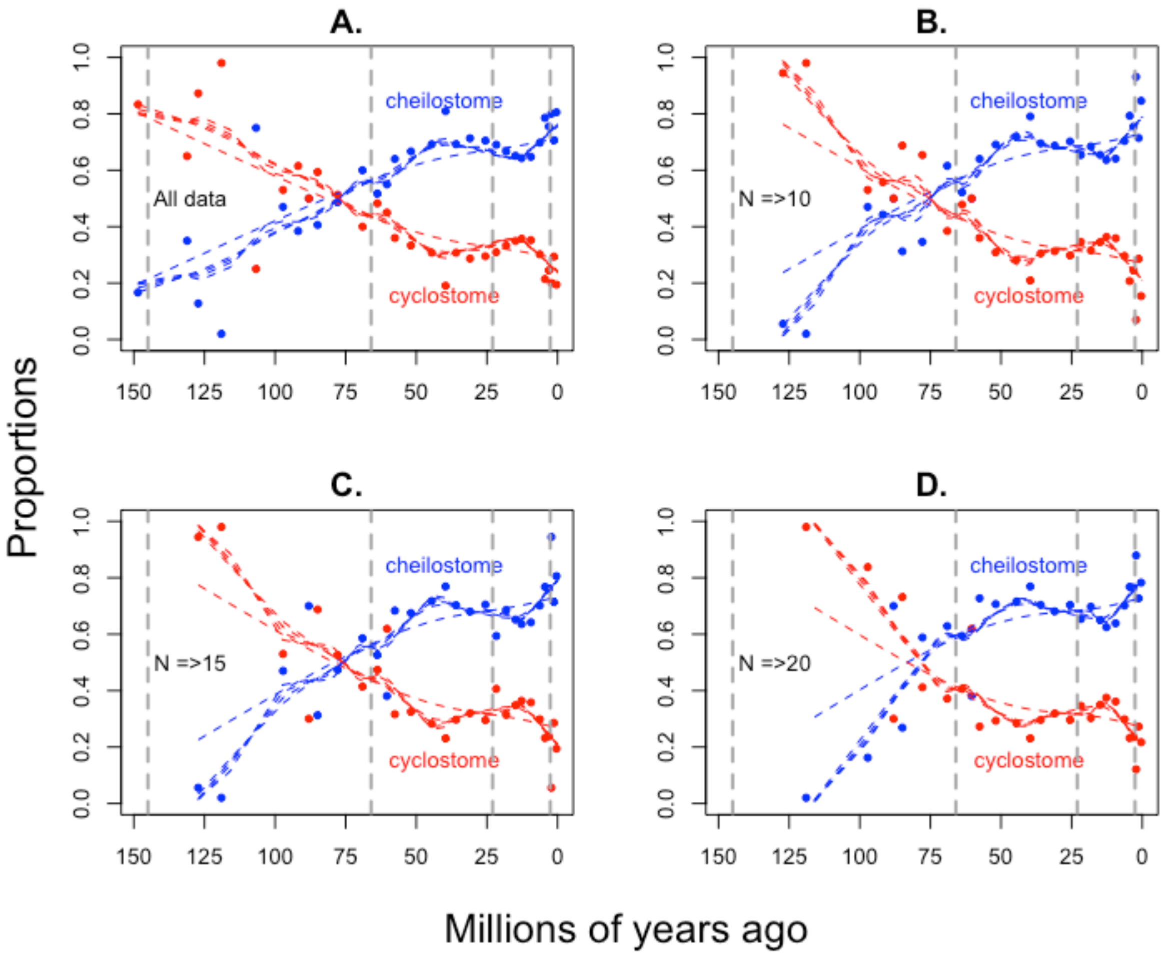

**fig. S3. Genus range-through curves.** A. This panel plots our new, synonymized genus data (solid circles and lines) for cheilostomes (blue) and cyclostomes (red) separately and together (black). The dashed lines (blue = cheilostomes, red = cyclostomes) show data from McKinney and Taylor (2001) which is the basis for even the most recent review, Taylor and Waeschenbach 2015. B. This panel repeats the synonymized genus data from A. for cheilostome and cyclostome genera separately but also plots the non-synonymized data (open circles and dotted lines) for comparison.

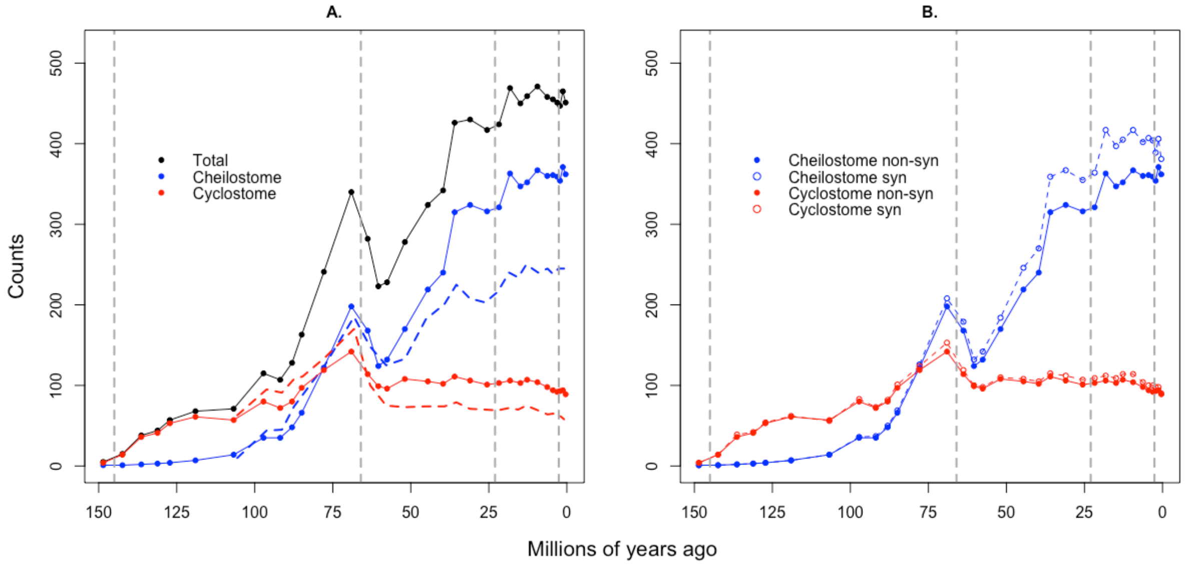

**fig. S4. Compare genus richness estimates from PyRate and Capture-recapture.** These panels compare A. cheilostome and B. cyclostome genus richness estimates from PyRate and a Jolly-Seber model, where 95% credibility and 95% confidence intervals are plotted, respectively. Estimates are plotted at the midpoint of time intervals but the PyRate estimates are shifted by 1 million year for visibility.

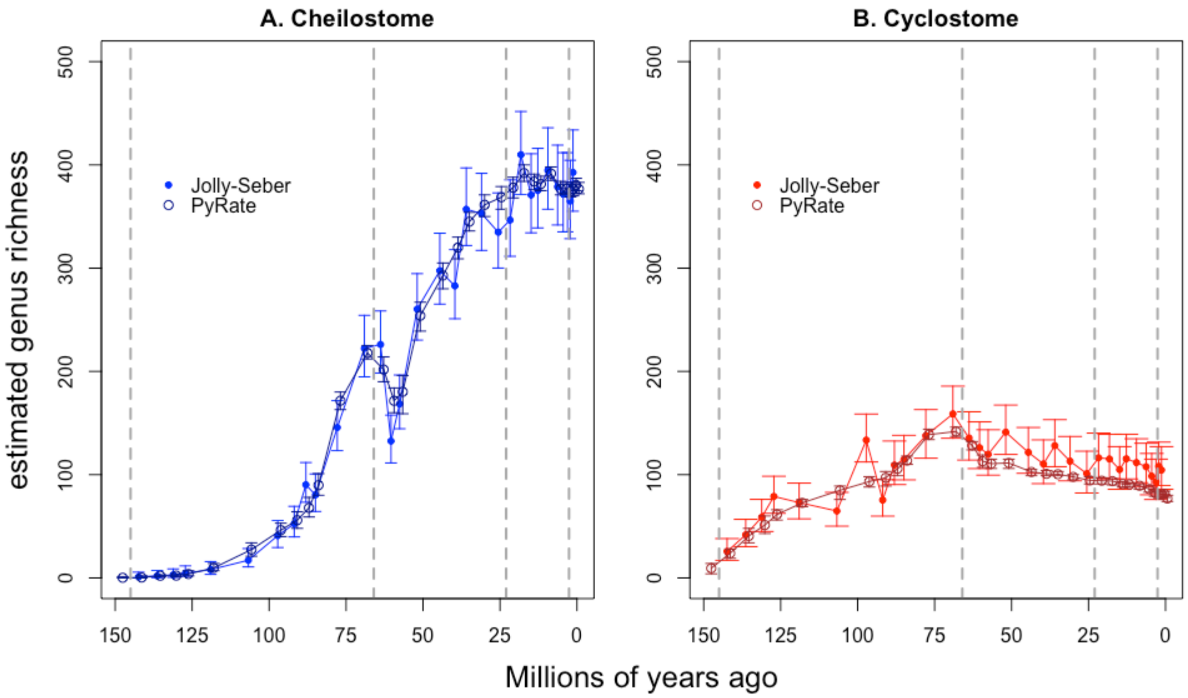

**fig. S5. Compare genus richness estimates with data that were manually collected to those compiled using automated text-mining.** These panels compare cheilostome and cyclostome genus richness estimates from a Jolly-Seber model using two data sources. The one labelled "manual" is the dataset of hand-collected, synonymized taxon occurrences presented in our main text, while the other, labelled "TMO" was compiled using Natural Language Processing (data from Kopperud et al. 2019). Bars represent 95% confidence intervals from the Jolly-Seber model.

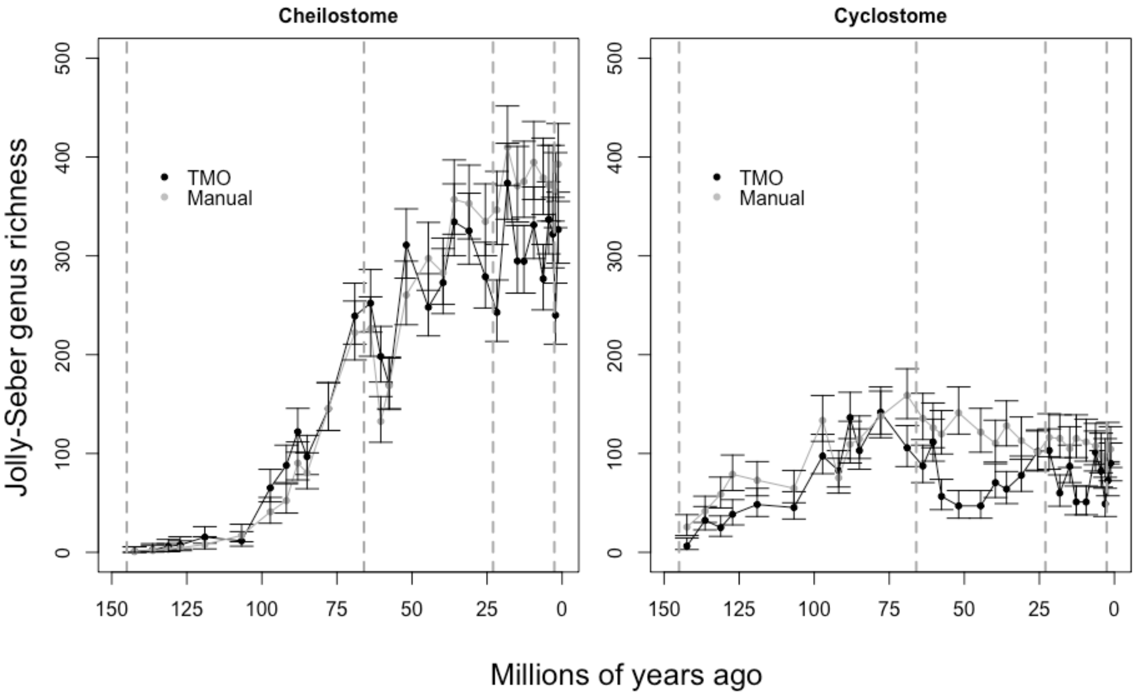

**fig. S6. Genus origination, extinction and sampling rates with confidence intervals.** Each panel shows the estimates from fully time-varying Pradel seniority models run separately for cheilostome genera (blue) and cyclostome genera (red). Only estimate that are relatively well-constrained are plotted (solid circles) and lines joining the estimates are for visual aid only. Vertical lines are 95% confidence intervals. A. Instantaneous origination rates B. Instantaneous extinction rates. C. Sampling rates. Note that sampling rates are for within internal stages while origination and extinction rates are for their boundaries and that the y-axes vary across the panels.

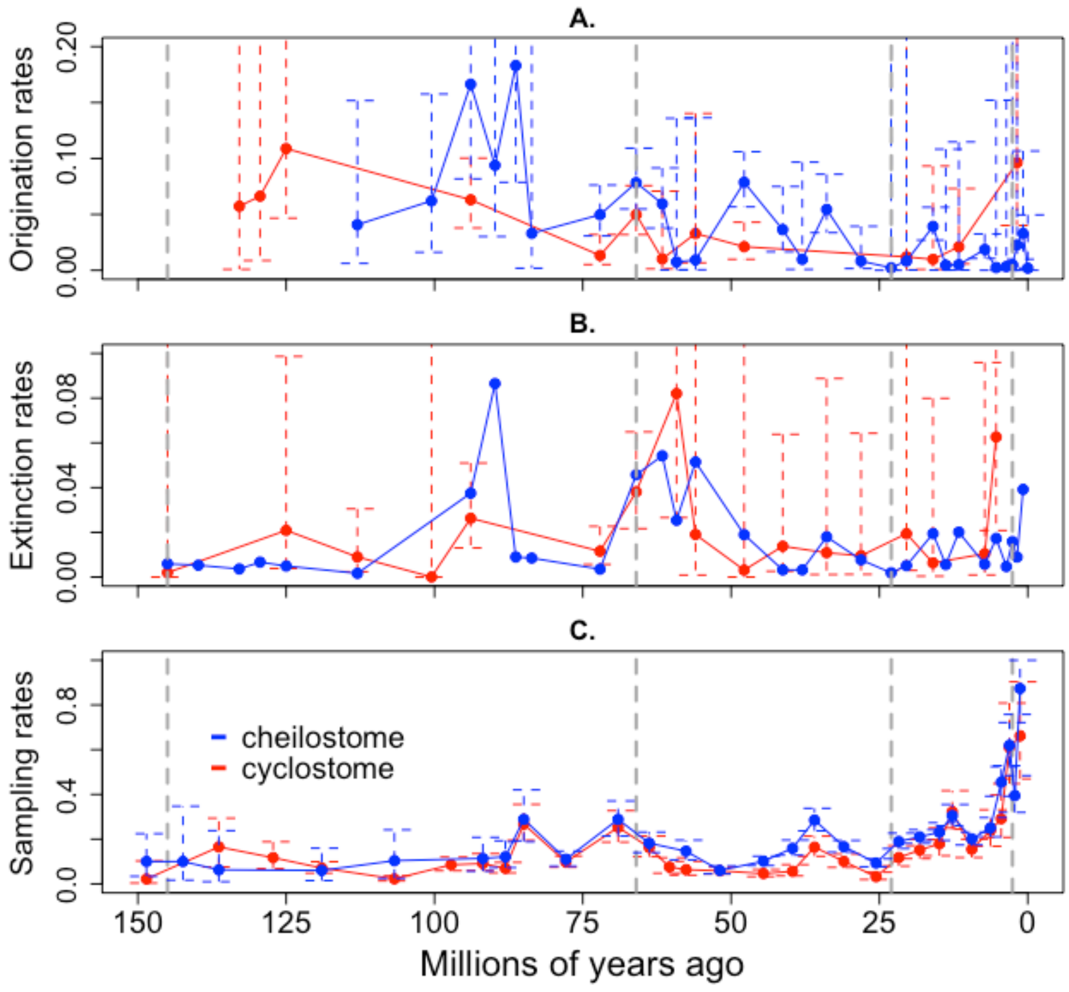

**fig. S7. Cyclostome genus origination, extinction and sampling rates: manual and text-mined data.** Each panel shows the estimates from fully time-varying Pradel seniority models run for cyclostome genera, comparing our combined data and text-mined data from Kopperud et al. 2019. Vertical lines are 95% confidence intervals from the combined data estimates. Note that confidence intervals could not be estimated for the text-mined data and are not plotted A. Instantaneous origination rates B. Instantaneous extinction rates. C. Sampling rates. Note that sampling rates are for within internal stages while origination and extinction rates are for their boundaries and that the y-axes vary across the panels.

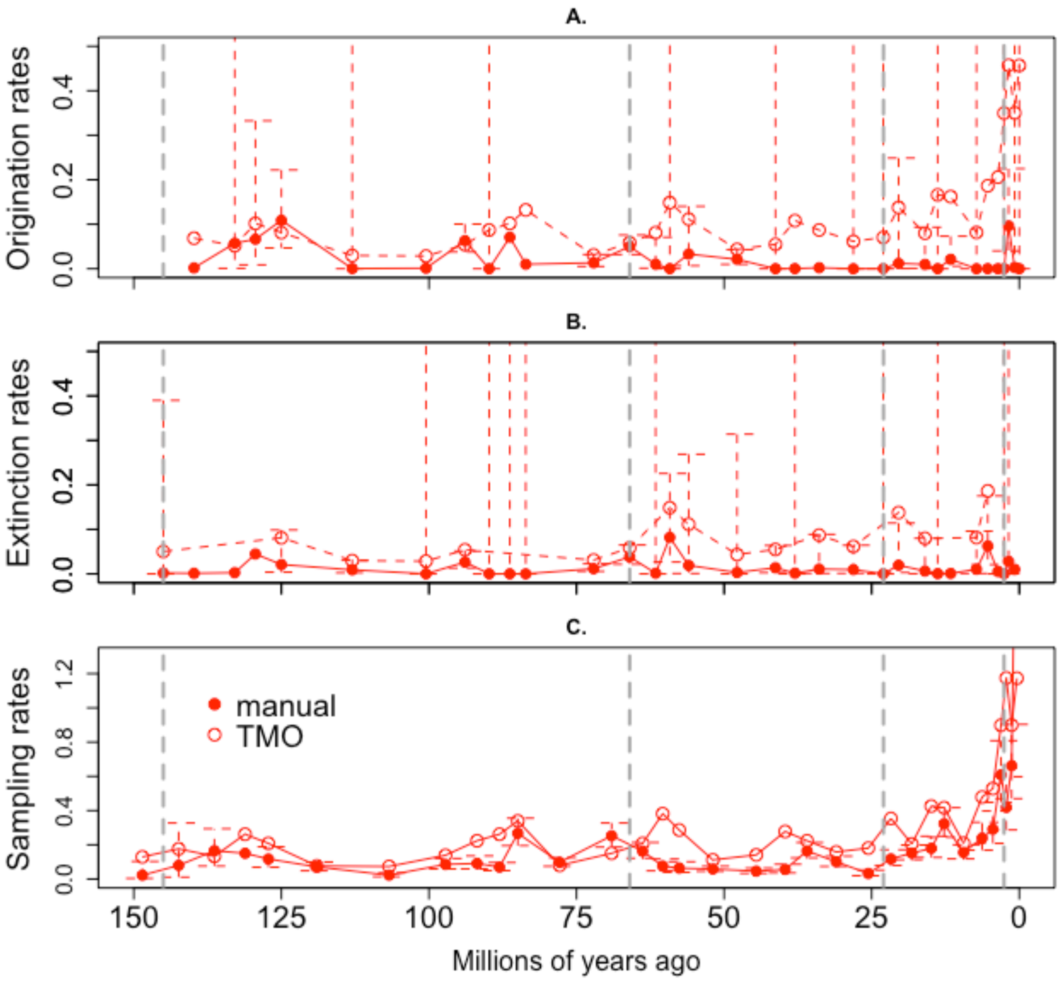

**fig. S8. Cheilostome genus origination, extinction and sampling rates: manual and text-mined data.** Each panel shows the estimates from fully time-varying Pradel seniority models run for cheilostome genera, comparing our combined data and text-mined data from Kopperud et al. 2019. Vertical blue lines are 95% confidence intervals from the combined data estimates. Vertical light blue lines are 95% confidence intervals from the TMO data estimates. TMO CI's are larger but overlap estimates from the combined dataset. A. Instantaneous origination rates B. Instantaneous extinction rates. C. Sampling rates. Note that sampling rates are for within internal stages while origination and extinction rates are for their boundaries.

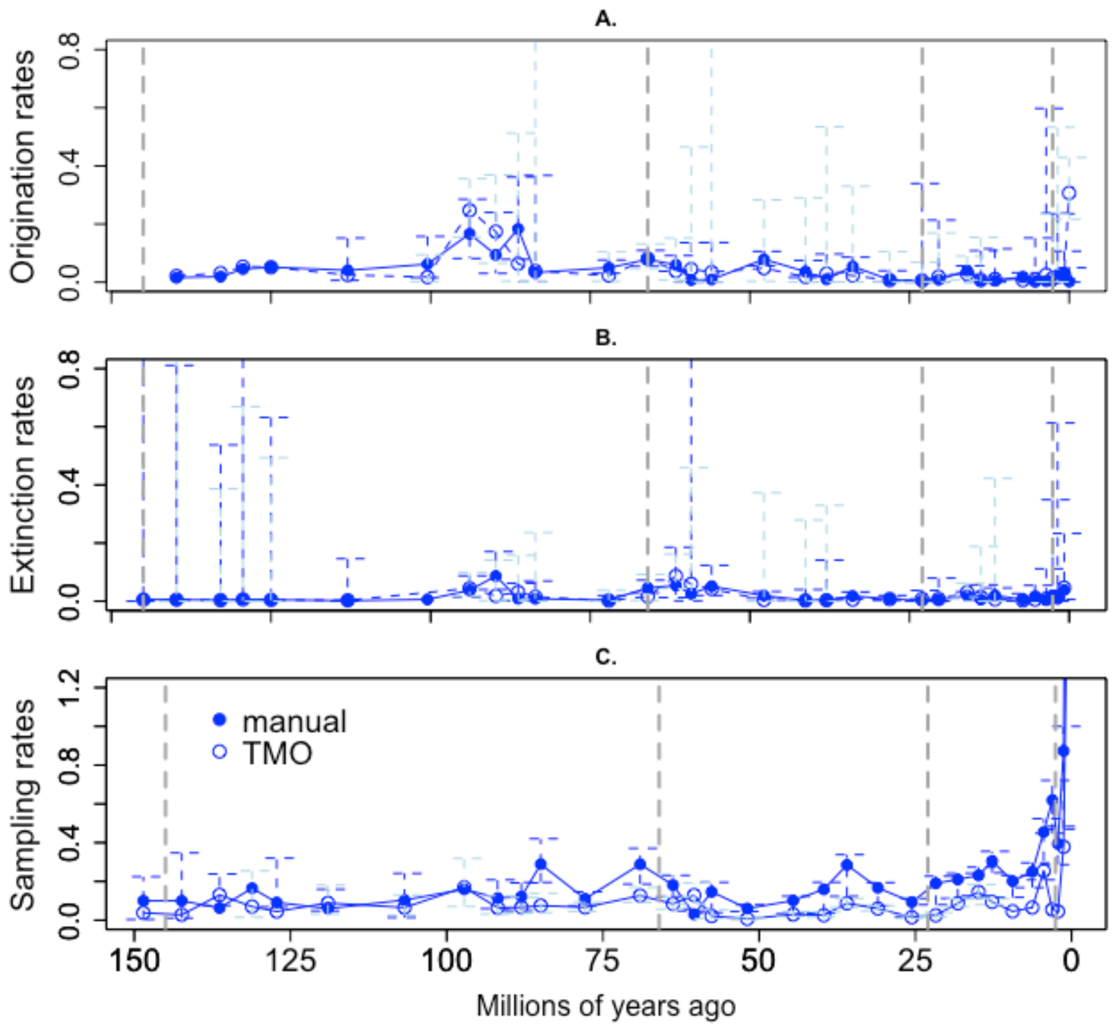

**fig. S9. Cyclostome genus origination & extinction (Pradel and PyRate)**

PyRate origination and extinction rates (black) are averaged from posterior MCMC outputs while Pradel estimates (red) are as in fig. S4. The same treatment is given to extinction rates.

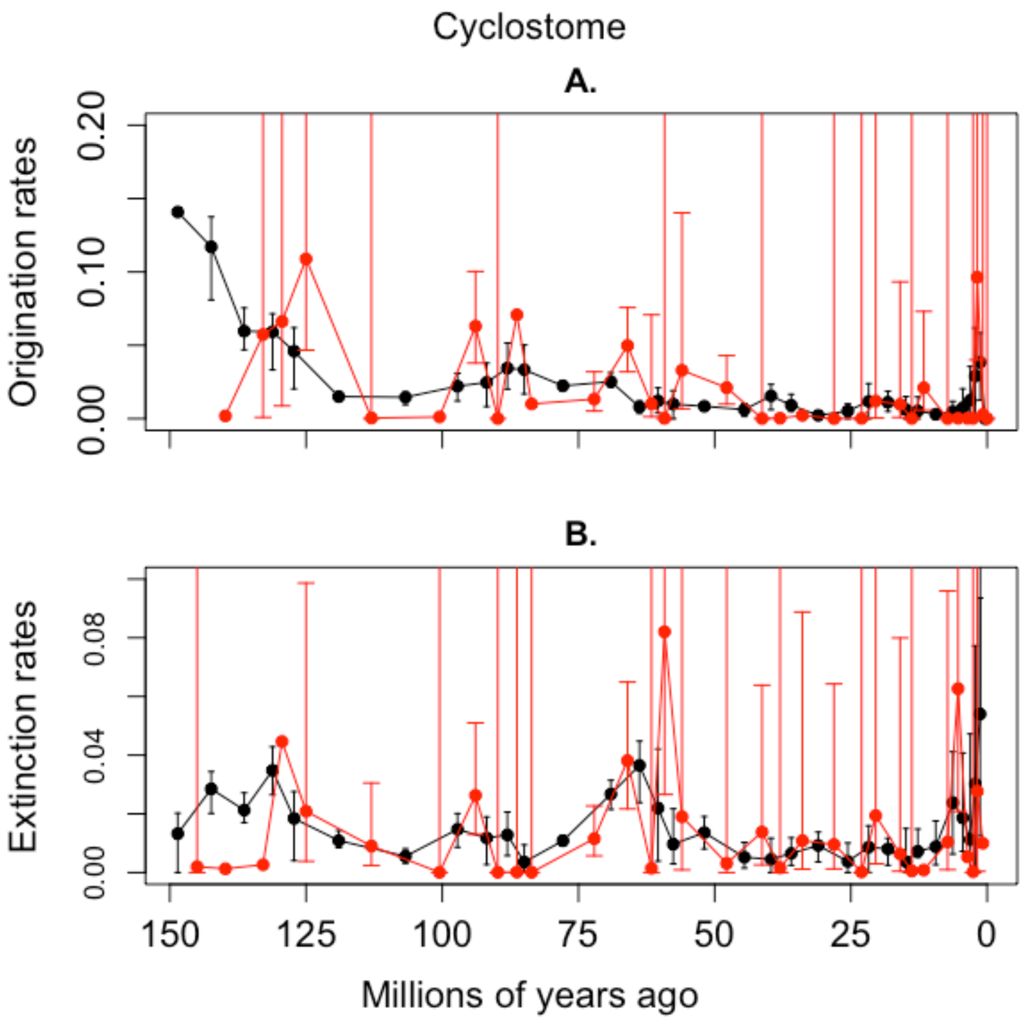

**fig. S10. Cheilostome genus origination & extinction (Pradel and PyRate)**

PyRate origination and extinction rates (black) are averaged from posterior MCMC outputs while Pradel estimates (blue) are as in fig. S4. The same treatment is given to extinction rates.

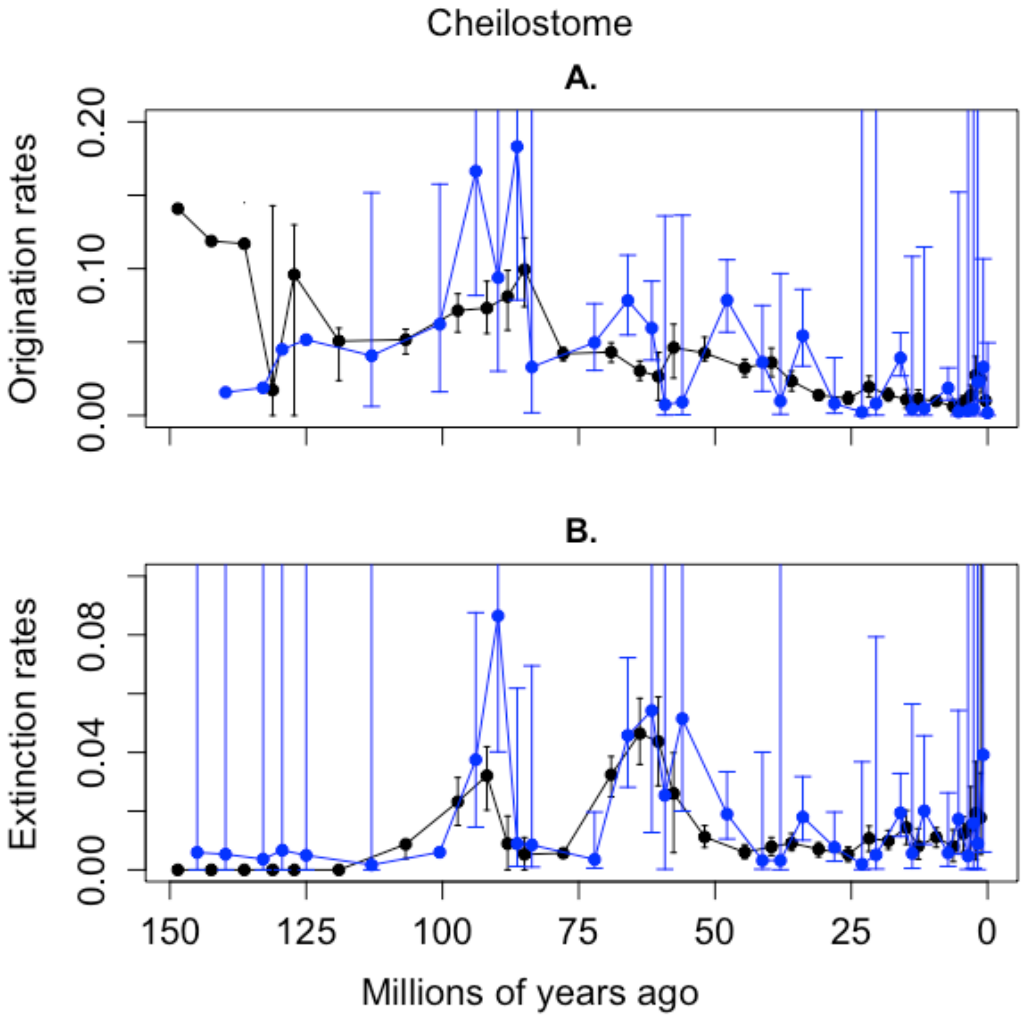

**Table S1. Relationship between Cheilostome and Cyclostome extinction.** Parameter estimates (mean, median and lower and upper bounds of the models shown in Table 1) are presented for the relationship between cheilostome and cyclostome extinction rates. The $\mu$s are means in the OU model, $dt$ are characteristic times, $\sigma$ stationary variances, and $\beta$s the strengths of the granger causal link between the two time series (see above and Reitan and Liow 2019 for details).

| **Model D** | Mean | Median | Lower 5% | Upper 95% |
|---|---|---|---|---|
| $\mu$.cheilostome.extinction | -2.126 | -2.088 | -3.641 | -0.605 |
| $dt$.cheilostome.extinction | 43.778 | 20.366 | 1.037 | 197.570 |
| $\sigma$.cheilostome.extinction | 0.288 | 0.294 | 0.002 | 0.623 |
| $\mu$.cyclostome.extinction | -2.459 | -2.527 | -3.728 | -0.939 |
| $dt$.cyclostome.extinction | 19.778 | 5.012 | 0.662 | 112.506 |
| $\sigma$.cyclostome.extinction | 0.195 | 0.152 | 0.002 | 0.6467 |
| $\beta$.cheilostome.extinction.to.cyclostome.extinction | 0.522 | 0.572 | -0.345 | 1.212 |
| $\beta$.cyclostome.extinction.to.cheilostome.extinction | 0.326 | 0.442 | -1.017 | 1.315 |
| **Model B** | | | | |
| $\mu$.cheilostome.extinction | -2.227 | -2.208 | -3.730 | -0.697 |
| $dt$.cheilostome.extinction | 72.621 | 52.952 | 6.592 | 272.045 |
| $\sigma$.cheilostome.extinction | 0.342 | 0.323 | 0.134 | 0.627 |
| $\mu$.cyclostome.extinction | -2.492 | -2.510 | -3.681 | -1.111 |
| $dt$.cyclostome.extinction | 7.916 | 3.401 | 0.462 | 43.597 |
| $\sigma$.cyclostome.extinction | 0.129 | 0.068 | 0.0008 | 0.490 |
| $\beta$.cheilostome.extinction.to.cyclostome.extinction | 0.679 | 0.693 | -0.007 | 1.278 |
| **Model C** | | | | |

| | | | | |
|---|---|---|---|---|
| $\mu$.cheilostome.extinction | -2.505 | -2.501 | -3.871 | -0.986 |
| $dt$.cheilostome.extinction | 13.332 | 3.924 | 0.650 | 90.172 |
| $\sigma$.cheilostome.extinction | 0.205 | 0.165 | 0.002 | 0.617 |
| $\mu$.cyclostome.extinction | -2.443 | -2.440 | -3.870 | -0.658 |
| $dt$.cyclostome.extinction | 65.501 | 42.013 | 2.472 | 250.230 |
| $\sigma$.cyclostome.extinction | 0.298 | 0.286 | 0.0072 | 0.647 |
| $\beta$.cyclostome.extinction.to.cheilostome.extinction | 0.702 | 0.776 | -0.493 | 1.439 |

**Table S2. Relationship between Cheilostome extinction and Cyclostome origination.** Parameter estimates (mean, median and lower and upper bounds of the models shown in Table 1) are presented for the relationship between cheilostome extinction and cyclostome origination rates. The $\mu$s are means in the OU model, $dt$ are characteristic times, $\sigma$ stationary variances, and $\beta$s the strengths of the Granger causal link between the two time series (see above and Reitan and Liow 2019 for details).

| **Model D** | Mean | Median | Lower 95% | Upper 95% |
|---|---|---|---|---|
| $\mu$.cheilostome.extinction | -2.482 | -2.493 | -3.795 | -1.070 |
| $dt$.cheilostome.extinction | 21.170 | 9.521 | 0.925 | 115.990 |
| $\sigma$.cheilostome.extinction | 0.197 | 0.168 | 0.002 | 0.577 |
| $\mu$.cyclostome. origination | -2.026 | -2.036 | -3.268 | -0.472 |
| $dt$.cyclostome. origination | 42.298 | 26.232 | 1.023 | 174.940 |
| $\sigma$.cyclostome.origination | 0.275 | 0.252 | 0.009 | 0.658 |
| $\beta$.cheilostome.extinction.to.cyclostome.origination | 0.275 | 0.303 | -0.915 | 1.185 |
| $\beta$.cyclostome.origination.to.cheilostome.extinction | 0.563 | 0.637 | -0.773 | 1.314 |
| **Model C** | | | | |
| $\mu$.cheilostome.extinction | -2.669 | -2.744 | -3.754 | -1.194 |
| $dt$.cheilostome.extinction | 23.605 | 9.255 | 0.893 | 158.506 |
| $\sigma$.cheilostome.extinction | 0.180 | 0.151 | 0.001 | 0.563 |
| $\mu$.cyclostome. origination | -2.234 | -2.278 | -3.326 | -0.689 |
| $dt$.cyclostome. origination | 57.193 | 38.156 | 2.445 | 248.063 |
| $\sigma$.cyclostome.origination | 0.292 | 0.268 | 0.083 | 0.657 |

| | | | | |
|---|---|---|---|---|
| $\beta$.cyclostome.origination.to.cheilostome.extinction | 0.621 | 0.673 | -0.425 | 1.345 |
| **Model B** | | | | |
| $\mu$.cheilostome.extinction | -2.750 | -2.821 | -4.053 | -0.947 |
| $dt$.cheilostome.extinction | 49.825 | 32.363 | 3.565 | 202.163 |
| $\sigma$.cheilostome.extinction | 0.297 | 0.275 | 0.094 | 0.649 |
| $\mu$.cyclostome. origination | -2.449 | -2.522 | -3.422 | -1.144 |
| $dt$.cyclostome. origination | 20.514 | 7.649 | 0.726 | 115.732 |
| $\sigma$.cyclostome.origination | 0.246 | 0.216 | 0.003 | 0.772 |
| $\beta$.cheilostome.extinction.to.cyclostome.origination | 0.350 | 0.406 | -0.801 | 1.202 |
